# Predicted genetic gains from introgressing chromosome segments from exotic germplasm into an elite soybean cultivar

**DOI:** 10.1101/701987

**Authors:** Sushan Ru, Rex Bernardo

**Affiliations:** University of Minnesota, Department of Agronomy and Plant Genetics, 411 Borlaug Hall, 1991 Upper Buford Circle, Saint Paul, MN 55108, USA

**Keywords:** targeted recombination, introgression, exotic germplasm

## Abstract

Broadening the diversity of cultivated soybean [*Glycine max* (L.) Merrill] through introgression of exotic germplasm has been difficult. Our objectives were to 1) determine if introgressing specific chromosome segments (instead of quantitative trait locus alleles) from exotic soybean germplasm has potential for improving an elite cultivar, and 2) identify strategies to introgress and pyramid exotic chromosome segments into an elite cultivar. We estimated genomewide marker effects for yield and other traits in seven crosses between the elite line IA3023 and seven soybean plant introductions (PIs). We then predicted genetic gains from having ≤2 targeted recombinations per linkage group. When introgression was modeled for yield while controlling maturity in the seven PI × IA3023 populations, the predicted yield was 8 to 25% over the yield of IA3023. Correlated changes in maturity, seed traits, lodging, and plant height were generally small but were in the favorable direction. In contrast, selecting the best recombinant inbred (without targeted recombination) in each of the PI × IA3023 populations led to negative or minimal yield gains over IA3023. In one PI × IA3023 population, introgressing and pyramiding only two linkage groups from recombinant inbreds into IA3023 was predicted to achieve an 8% yield gain over IA3023 without sacrificing the performance of other traits. The probability of inheriting intact chromosomes was high enough to allow introgression and pyramiding of chromosome segments in 5-6 generations. Overall, our study suggested that introgressing specific chromosome segments is an effective way to introduce exotic soybean germplasm into an elite cultivar.

**Key message:** To improve an elite soybean line, introgress longer chromosome segments instead of QTL alleles from exotic germplasm.

## Introduction

Cultivated soybean [*Glycine max* (L.) Merrill] has limited diversity (Hyten et al. 2006), with only 17 founder lines accounting for >84% of the genetic base of modern soybean cultivars (Gizlice et al. 1994). This lack of diversity in elite soybean germplasm hampers future genetic gains (St. Martin 1982, Delannay et al. 1983, Cooper et al. 2001). To date, soybean breeders have used two main approaches to introgress exotic germplasm or genes in an attempt to broaden the soybean genetic base.

First, soybean breeders have made exotic × elite crosses with the goal of finding superior transgressive segregants among F_2_- or backcross-derived lines (Carter et al. 2004). This approach has generally been unsuccessful because soybean plant introductions (PIs) tend to have low yield and excessive shattering and lodging (Sneller et al. 1997, Carter et al. 2004). This low mean performance of soybean PIs has led to a low probability of finding progeny superior for multiple traits (Thorne and Fehr 1970b, Thorne and Fehr 1970a, Vello et al. 1984, Ininda et al. 1996). These empirical results are supported by theoretical studies showing that when the two parents differ substantially in performance, the probability of finding a line superior to the better parent is often low (Bailey and Comstock 1976).

Second, soybean breeders have attempted to detect and introgress quantitative trait locus (QTL) alleles found in exotic germplasm but absent in elite germplasm. The rationale is to introgress only a selected set of very small chromosome segments (i.e., marker intervals that harbor QTL) from exotic germplasm to elite germplasm, thereby avoiding the decrease in the population mean with exotic × elite crosses in the first approach. However, this second approach has been largely unsuccessful for yield because detectable QTL alleles for soybean yield have been few (Concibido et al. 2003, Diers et al. 2018), their effects tended to be small (Diers et al. 2018), and their effects were often inconsistent across genetic backgrounds (Concibido et al. 2003). Furthermore, QTL alleles for yield often have a negative influence on other traits such as lodging and maturity (Guzman et al. 2007, Kim et al. 2012).

Instead of introgressing individual QTL alleles, an alternative approach is to introgress longer chromosome segments that harbor favorable alleles found in exotic soybean germplasm. Suppose linkage group (LG) 1 has single nucleotide polymorphism (SNP) markers 1, 2, 3, …, 20. An analysis of genomewide marker effects (Bernardo 2017, Brandariz and Bernardo 2018, Ru and Bernardo 2019) may indicate that, for example, introgressing the chromosome segment between SNP 5 and SNP 10 would improve the performance of the elite line for multiple traits. This targeted introgression of chromosome segments is likely superior to QTL introgression because the six-SNP chromosome segment in this example could capture the effects of several minor QTLs, rather than the effect of a single minor QTL. Introgression of specific chromosome segments would be an application of targeted recombination, which refers to inducing or selecting for specific recombination points on chromosomes to increase genetic gains in a cross (Bernardo 2017). Targeted recombination was found to double the predicted gains for yield and other agronomic traits in maize (*Zea mays* L.), soybean, pea (*Pisum sativum* L.), wheat (*Triticum aestivum* L.), and barley (*Hordeum vulgare* L.) (Bernardo 2017, Brandariz and Bernardo 2018, Ru and Bernardo 2019).

The usefulness of introgressing specific chromosome segments from soybean PIs into an elite line has not yet been reported. Our objectives were to 1) determine if introgressing specific chromosome segments from exotic soybean germplasm has potential for improving an elite cultivar, and 2) identify feasible strategies to backcross and pyramid natural recombinations into elite cultivars.

## Materials & Methods

### Populations, phenotypic, marker, and linkage mapping data

We studied eight biparental populations from the soybean nested association mapping (NAM) study, which had the elite line IA3023 as the common, elite parent (Diers et al. 2018) and included seven PI lines and a high-yielding line (4J105-3-4) as unique parents (Table 1). The elite × elite cross between IA3023 and 4J105-3-4 was used for comparison. The PI parents had much lower yield than IA3023 but performed well under severe drought (Diers et al. 2018). Up to 140 recombinant inbreds were available in each cross. Phenotypic and SNP marker data on the Wm82.a2 soybean assembly were downloaded from SoyBase (www.soybase.org/SoyNAM/, accessed October 30, 2018). Phenotypic data used in this study included yield (kg/ha), 100-seed weight (g), protein content (g/kg), oil content (g/kg), lodging (1-5 scale with 5 being the most lodging), plant height (cm), and maturity (d) across 22 environments in the United States from 2011 to 2013 (Diers et al. 2018). The linkage map for each cross was obtained from Song et al. (2017).

**Table 1.** Soybean nested association mapping (NAM) populations used in this study

| **Population** | **Cross** | **Lines** | **Markers** | **LG<sup>a</sup>** |
| --- | --- | --- | --- | --- |
| NAM03 | IA3023 × 4J105-3-4 | 137 | 890 | 33 |
| NAM40 | IA3023 × PI 398881 | 137 | 1166 | 33 |
| NAM41 | IA3023 × PI 427136 | 99 | 925 | 35 |
| NAM42 | IA3023 × PI 437169B | 133 | 1773 | 27 |
| NAM48 | IA3023 × PI 518751 | 139 | 1669 | 32 |
| NAM50 | IA3023 × PI 561370 | 132 | 1338 | 29 |
| NAM54 | IA3023 × PI 404188A | 140 | 1454 | 36 |
| NAM64 | IA3023 × PI 574486 | 124 | 1185 | 31 |

<sup>a</sup> linkage group

### Estimation of genomewide marker effects

Raw SNP data were filtered and imputed prior to estimating genomewide marker effects. First, markers that had the same genomic position were filtered so that only one marker was used at any unique linkage map position. Second, markers with >50% of missing values were removed, after which LGs with ≤5 markers were also disregarded. Missing genotypes on the remaining markers were imputed based on the genotypes of adjacent markers and recombination frequency between the imputed marker and its adjacent markers (Jacobson et al. 2015). Recombination frequency between adjacent markers was converted from Kosambi map distance according to Lynch and Walsh (1998).

Genomewide marker effects were estimated for each of the eight crosses by ridge regression-best linear unbiased prediction, as implemented in rrBLUP (Endelman 2011, version 4.5, available at https://cran.r-project.org/src/contrib/Archive/rrBLUP/, accessed October 30, 2017). Environments were treated as having fixed effects in the mix.solve function. Predictive ability was calculated as the correlation between marker-predicted and observed performance (*r*_MP_) for each trait.

### Modeling targeted recombination

We considered selection for high yield, 100-seed weight, protein content, and oil content and for low lodging score. We first modeled targeted recombination for each trait independently according to three steps (Ru and Bernardo 2019). First, we considered all possible gametes with 0, 1, or 2 recombination events for each LG, based on the genotype of the F_1_ in each cross. Second, we identified the best doubled haploid genotype for each LG based on sum of effects of the marker alleles carried by the doubled homologue, i.e., 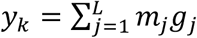, where *y*_*k*_ was the predicted performance at LG *k*; *m*_*j*_ was the estimated marker effect at locus *j*; ***g***_j_ was the genotype (−1 for one homozygote and 1 for the other homozygote) at locus *j*, and *L* was the total number of marker loci on the LG. Third, the predicted performance from targeted recombination across all LGs was calculated from the best *y*_*k*_ values as 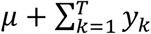, where *μ* was the mean performance of the cross as estimated in rrBLUP (Endelman 2011, version 4.5) and *T* was the total number of LGs.

Genetic gain from targeted recombination (*G*_Targeted_) was calculated as the marker-predicted performance of the ideal genotype with targeted recombination minus the marker-predicted performance of the elite parent, IA3023. Genetic gain from nontargeted recombination (*G*_Nontargeted_) was calculated as the marker-predicted performance of the best recombinant inbred found in the population minus the marker-predicted performance of IA3023. The significance of the increase in genetic gain from targeted recombination compared to nontargeted recombination (*G*_Targeted_ - *G*_Nontargeted_ >0) and the significance of *G*_Targeted_ >0 were tested by bootstrapping as described by Ru and Bernardo (2019).

To prevent unwanted changes in maturity, we also identified targeted recombination positions to maximize yield gain while holding maturity constant. For each LG, we identified doubled haploid genotypes that had a predicted maturity effect less than or equal to that of IA3023 on the same LG. We subsequently identified the one with the highest predicted yield. Similarly, we identified the highest yielding lines with a predicted maturity less than or equal to that of IA3023 for nontargeted recombination. The values of *G*_Targeted_ and *G*_Nontargeted_ were calculated the same way as in single-trait selection.

### Identifying recombinant inbreds with the most favorable recombinations for introgression

The 99–140 recombinant inbreds developed within each of the NAM populations (Table 1) can already serve as donor lines to improve IA3023. We therefore identified existing recombinant inbreds that had LGs that could be backcrossed and pyramided into IA3023. First, we calculated *y*_*k*_ for each LG in each recombinant inbred and identified the highest *y*_*k*_ for yield, with and without controlling maturity. Second, we determined whether any of the LGs among the existing recombinant inbreds had the 0–2 targeted recombinations that we had modeled. Third, we considered a NAM population as suitable for introgression if the predicted gain from pyramiding ≤4 LGs was greater than 170 kg/ha, which was the nontargeted yield gain with controlled maturity in the biparental cross between elite parents (NAM03).

We then designed strategies to pyramid the most favorable natural recombination events into the background of IA3023. Desirable yield gain with controlled maturity can be achieved in two ways. One way is to backcross and pyramid LGs with higher *y*_*k*_ for yield and equal or lower *y*_*k*_ for maturity relative to IA3023. The second way is to select LGs with complementing features, some to increase yield and others to reduce maturity.

Because a donor line already carries favorable recombinations on a specific LG, this LG needs to be passed intact from parent to offspring. We first calculated the probability of inheriting an intact LG from a parent during one round of meiosis in the following manner. Assuming the frequency of having at least one recombination on a LG is *p* and the frequency of having no recombination on a LG is *q* = 1 – *p*, we derived the probability of observing nonrecombinant genotypes for a given LG in the F5 generation, which corresponded to the recombinant-inbred generation in the soybean NAM populations (S1). We then calculated the expected ratio of nonrecombinant genotypes in the F5 (*r*_*p*_) for any *q* ranging from 0 to 1 with increments of 0.001. We subsequently compared the observed *r*_*p*_ with the derived *r*_*p*_ to identify the *p* and *q* values that result in a *r*_*p*_ value closest to the observed *r*_*p*_. Because only one of the two homologues is favorable, the probability of inheriting an intact LG from the parent with targeted recombination is *q*/2.

## Results

### Predicted gains from targeted recombination for single traits and multiple traits

Predicted performance of individual lines based on genomewide marker effects was highly correlated with observed performance for all traits in all crosses (*r*_MP_ > 0.8, Table 1). The common parent, IA3023, was superior to each PI parent for yield, lodging, and oil content, whereas each PI parent outperformed IA3023 for 100-seed weight and protein content (Table 2). The PI parents also tended to be much earlier maturing and taller than IA3023. In contrast, the high-yielding parent 4J105-3-4 (NAM03) was superior to IA3023 for yield, 100-seed weight, and protein content, but it had lower oil content.

**Table 2.**
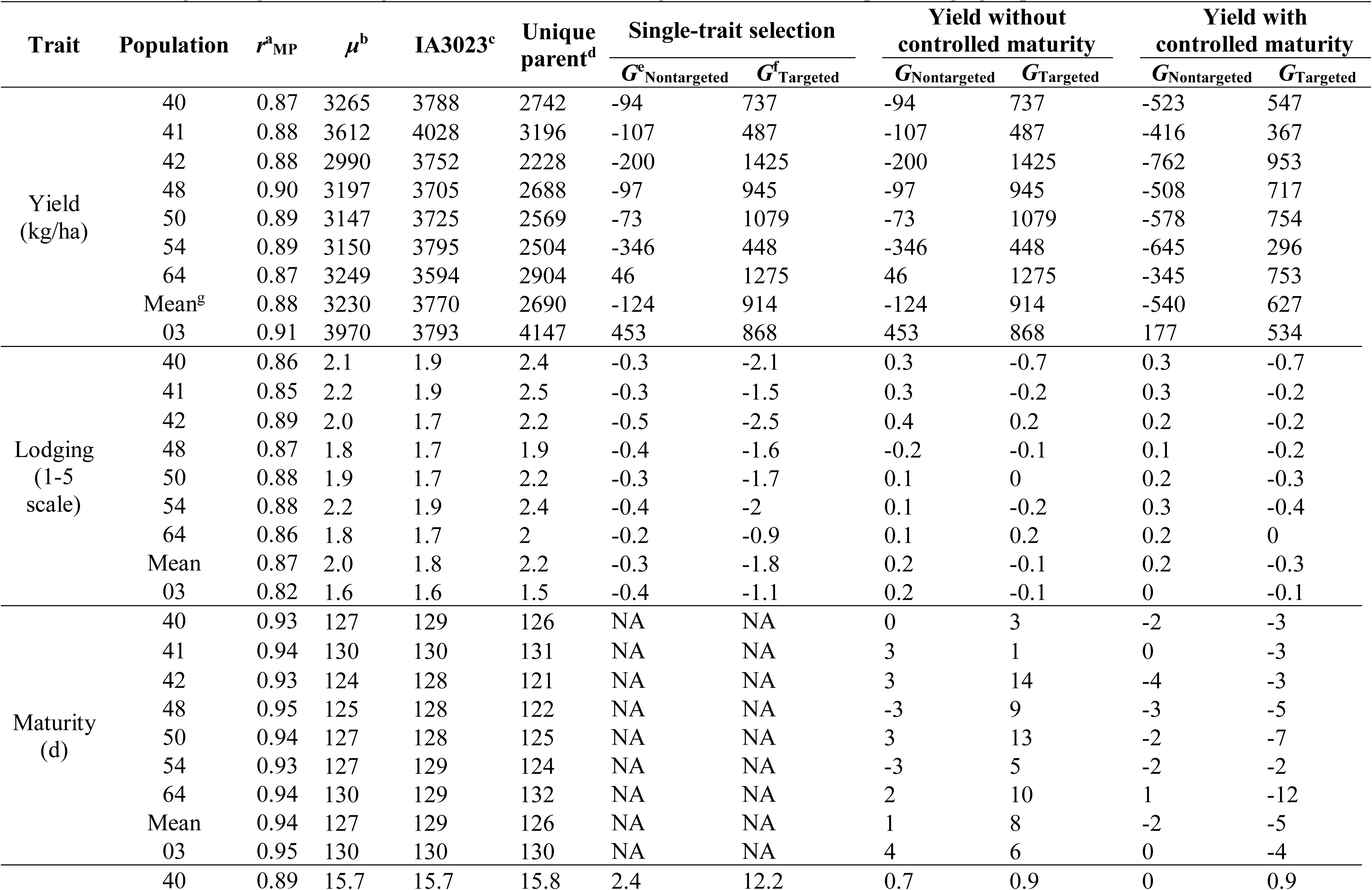
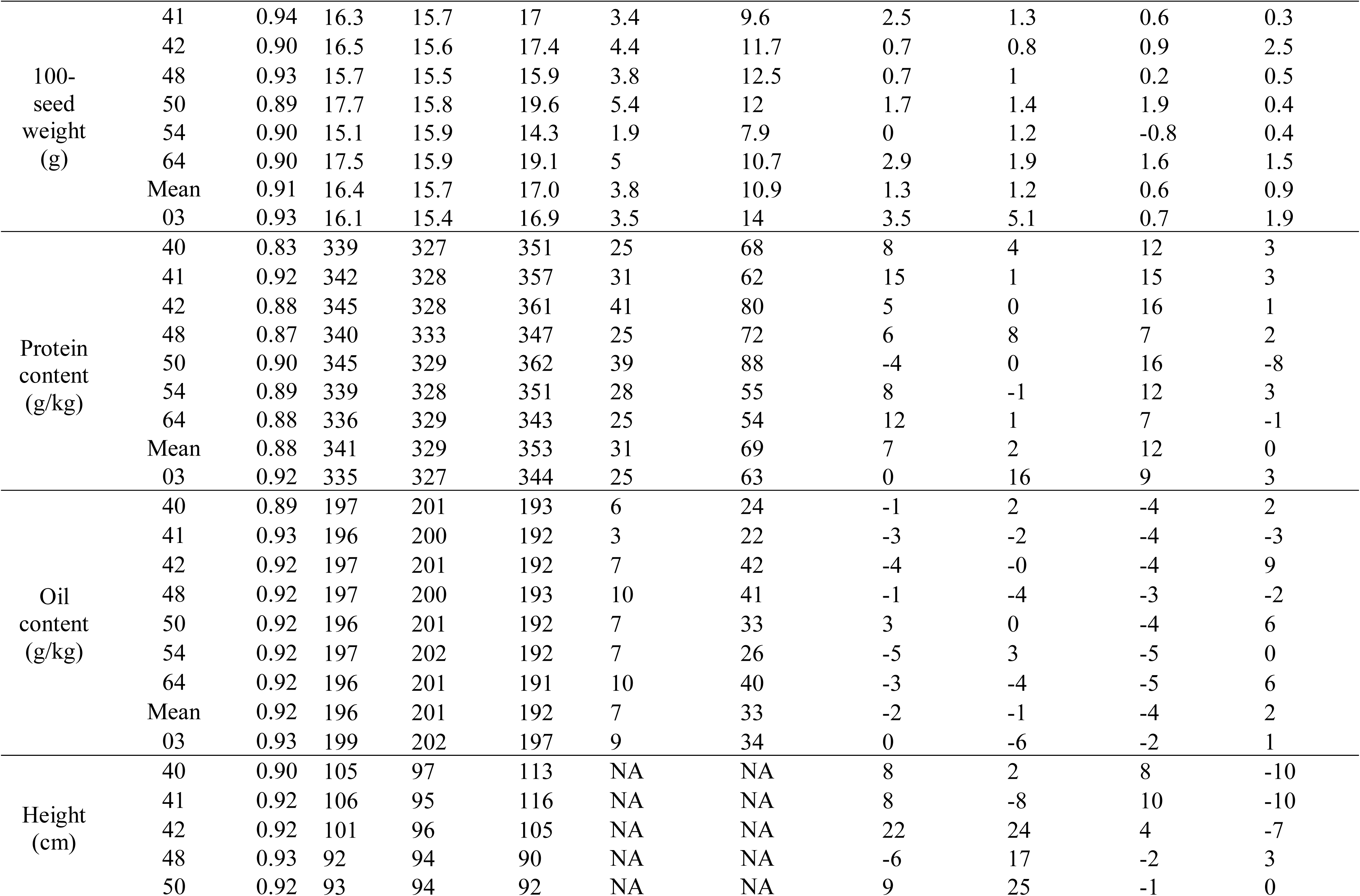
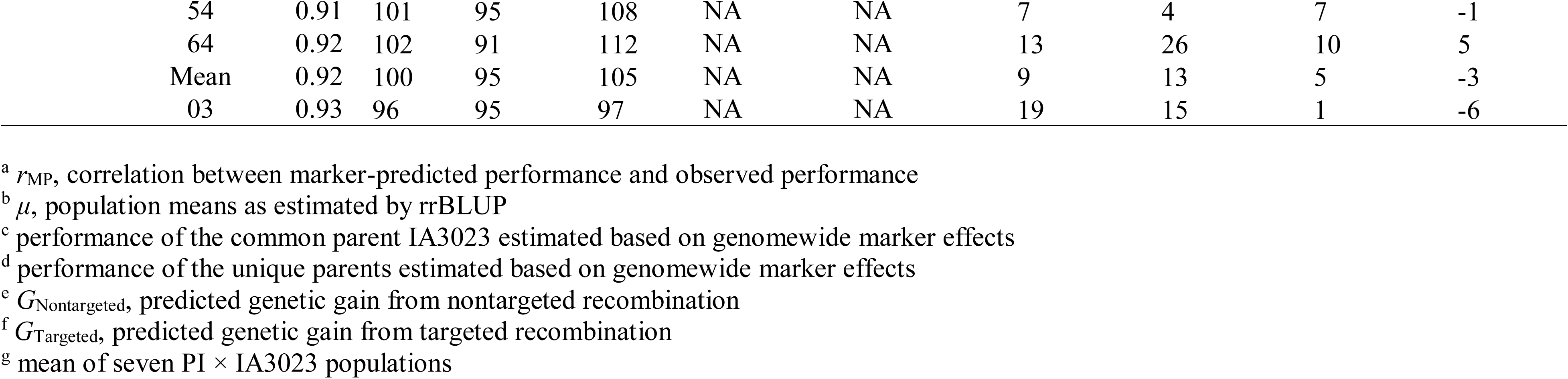
Predicted genetic gain from targeted recombination assuming 0-2 recombinations per linkage group

The highest-yielding lines in six out of seven PI × IA3023 populations were lower yielding than IA3023, whereas the best line in NAM03 had 12% higher yield (gain of 453 kg/ha) than IA3023 (Table 2). But with targeted recombination, the lines derived from the PI × IA3023 populations were predicted to significantly (*P* < 0.05) outyield IA3023 by 12-38% (Table 2). The predicted yield gains were higher in four out of the seven PI populations than in NAM03 (Table 2). On the other hand, targeted recombination to increase yield increased the predicted maturity (relative to IA3023) by 1 to 14 d (1-11%).

With the performance of IA3023 as the baseline, individual-trait selection without targeted recombination led to predicted gains of 10-28% for reduced lodging, 12-34% for 100-seed weight, 8-13% for protein content, and 1-5% for oil content in the PI × IA3023 populations and in NAM03 (Table 2). Compared with nontargeted recombination, targeted recombination in the PI × IA3023 populations led to significantly (*P* < 0.05) higher predicted gains of 51-143% for reduced lodging, 50-81% for 100 seed weight, 16-27% for protein content, and 11-21% for oil content.

When targeted recombination was modeled to control maturity, the resulting lines were predicted to have 1-9 d shorter maturity than IA3023 across all populations. This shorter maturity was accompanied by a significant (*P <* 0.05) increase in predicted yield (8-25% over the yield of IA3023), compared to a 14% yield gain in NAM03 (Table 2). With controlled maturity, predicted gains for yield were higher in five out of seven PI × IA3023 populations than in NAM03 (Table 2). Selecting for yield with controlled maturity led to a predicted decrease of 0.3 (16%) in lodging and small but positive changes (0-6%) in 100-seed weight, protein content, oil content, and plant height relative to IA3023 (Table 2).

### Strategies to introgress exotic chromosome segments into elite cultivar

Among the 191 LGs across eight populations where 1-2 targeted recombinations were modeled for desirable yield and maturity, 27 (14%) of the modeled LG genotypes were found present among the existing recombinant inbreds. For 13 of these LGs, the predicted yield was higher with natural targeted recombinations than with the 1-2 modeled recombinations because more than two natural recombinations occurred in a particular recombinant inbred. Among the donor lines, 14 therefore carried the 1-2 recombinations that were modeled and 13 carried natural recombinations that were not previously modeled but were predicted to have superior yield and controlled maturity.

We found that higher yield and similar maturity compared with IA3023 can be achieved if ≤4 LGs from selected recombinant inbred in each PI × IA3023 population are introgressed and pyramided into IA3023 (Table 3). Predicted gains from pyramiding ≤ 4 LGs in NAM42, NAM50, NAM40, NAM48, NAM41, and NAM64 were even higher than the predicted gain from selecting the highest-yielding inbred with ideal maturity in NAM03. Yield gain with negative or zero maturity change can be achieved in two ways. In NAM64, for example, one way is to backcross and pyramid three LGs, each controlling maturity, into the background of IA3023. This method had a predicted yield gain of 178 kg/ha (5% gain over IA3023). The second way is to pyramid LG3.1 in RIL DS11.64126 for its high yield effect (gain of 273 kg/ha) despite its large maturity effect (delay of 2 d) and LG10.3 from RIL DS11.64111 to reduce maturity by 3 d and slightly increase yield by 1 kg/ha (Fig. 1). Such introgression had predicted gains of 274 kg/ha for yield (8% gain over IA3023) and no change in maturity. Predicted changes in the six other traits were favorable but small (0-6%).

**Table 3.** Predicted yield gain (kg/ha) from introgressing and pyramiding ≤4 linkage groups from recombinant inbreds into IA3023

| Population | No. LG<sup>a</sup> | Yield<sup>d</sup> |  | Yield with controlled maturity<sup>e</sup> |  |
| --- | --- | --- | --- | --- | --- |
| | | $G^b_{\text{yield}}$ (kg/ha) | $G^c_{\text{maturity}}$ (d) | $G_{\text{yield}}$ (kg/ha) | $G_{\text{maturity}}$ (d) |
| 42 | 1 | 153 | 0 | 120 | -1 |
|  | 2 | 261 | 1 | 226 | -1 |
|  | 3 | 368 | 1 | 297 | -1 |
|  | 4 | 455 | 2 | 340 | -2 |
| 54 | 1 | 53 | 1 | 37 | 0 |
|  | 2 | 95 | 1 | 74 | -1 |
|  | 3 | 132 | 1 | 105 | -1 |
|  | 4 | 169 | 1 | 133 | -1 |
| 50 | 1 | 114 | 3 | 58 | 0 |
|  | 2 | 201 | 6 | 115 | 0 |
|  | 3 | 276 | 9 | 172 | -1 |
|  | 4 | 344 | 10 | 224 | -1 |
| 40 | 1 | 65 | 0 | 54 | 0 |
|  | 2 | 121 | 1 | 103 | -1 |
|  | 3 | 174 | 1 | 147 | -1 |
|  | 4 | 226 | 1 | 189 | -1 |
| 48 | 1 | 128 | 0 | 124 | 0 |
|  | 2 | 221 | 1 | 199 | 0 |
|  | 3 | 308 | 2 | 261 | 0 |
|  | 4 | 389 | 2 | 315 | 0 |
| 41 | 1 | 65 | 0 | 46 | 0 |
|  | 2 | 128 | 1 | 87 | -1 |
|  | 3 | 170 | 1 | 126 | -1 |
|  | 4 | 209 | 0 | 161 | -2 |
| 64 | 1 | 272 | 2 | 69 | -0 |
|  | 2 | 401 | 5 | 125 | -1 |
|  | 3 | 475 | 5 | 178 | -2 |
|  | 4 | 539 | 6 | 223 | -3 |
| 03 | 1 | 160 | 1 | 94 | -0 |
|  | 2 | 236 | 4 | 150 | -0 |
|  | 3 | 307 | 5 | 199 | -0 |
|  | 4 | 377 | 5 | 243 | -1 |

<sup>a</sup> number of linkage groups to introgress and pyramid into IA3023
<sup>b</sup> $G_{\text{yield}}$ , maximum yield gain from pyramiding a given number of linkage groups
<sup>c</sup> $G_{\text{maturity}}$ , estimated maturity gain when the maximum yield gain was achieved from pyramiding a given number of linkage groups
<sup>d</sup> linkage groups were selected based on the increase in yield over IA3023
<sup>e</sup> linkage groups were selected for higher yield and lower maturity relative to IA3023

**Fig. 1.**
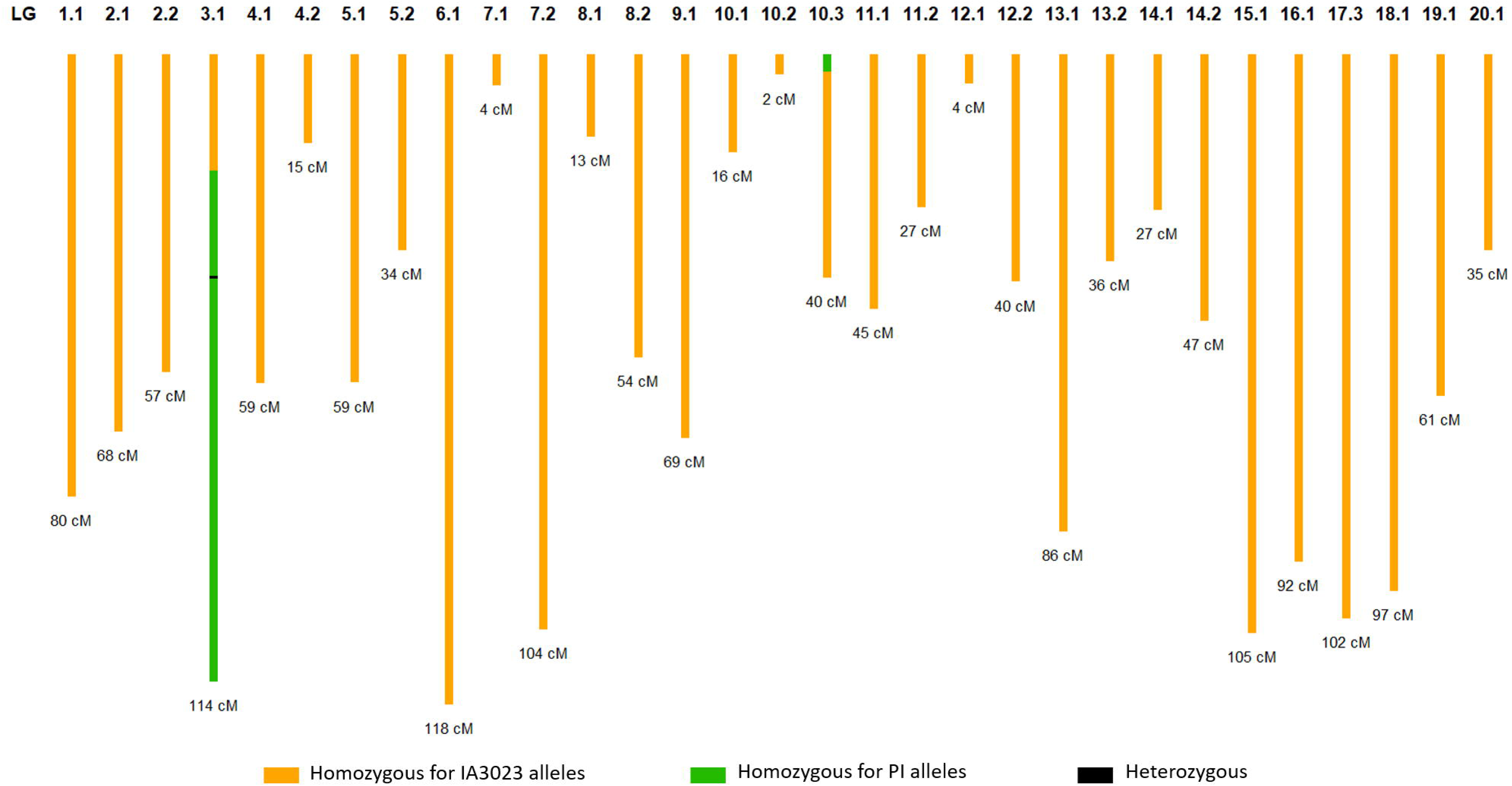
Targeted genotype for introgressing LGs 3.1 and 10.3 from NAM64 inbreds into IA3023.

Based on the genotypic data in NAM64, the estimated frequency of parental genotypes among recombinant inbreds was *r*_*p*_ = 4% for LG3.1 and 43% for LG10.3. The estimated probability of inheriting the desired, intact chromosome during one round of backcrossing was *q*/2 = 12.5% for LG3.1 and *q*/2 = 35% for LG10.3. The estimated *q* values indicated a two-step procedure to pyramid targeted recombinations on LG3.1 and LG10.3 into the background of IA3023. First, DS11.64126 and DS11.64111 can be backcrossed independently to the recurrent parent IA3023 to obtain offspring with LG3.1 from DA11.64126 and offspring with LG10.3 from DS11.64111 (Fig. 2). Second, once the targeted recombination from LG3.1 and LG10.3 has been introduced and the rest of the genome is fixed with alleles from IA3023, selected offspring from two independent backcross lines can be crossed and selfed to pyramid targeted recombinations from both LGs.

**Fig. 2.**
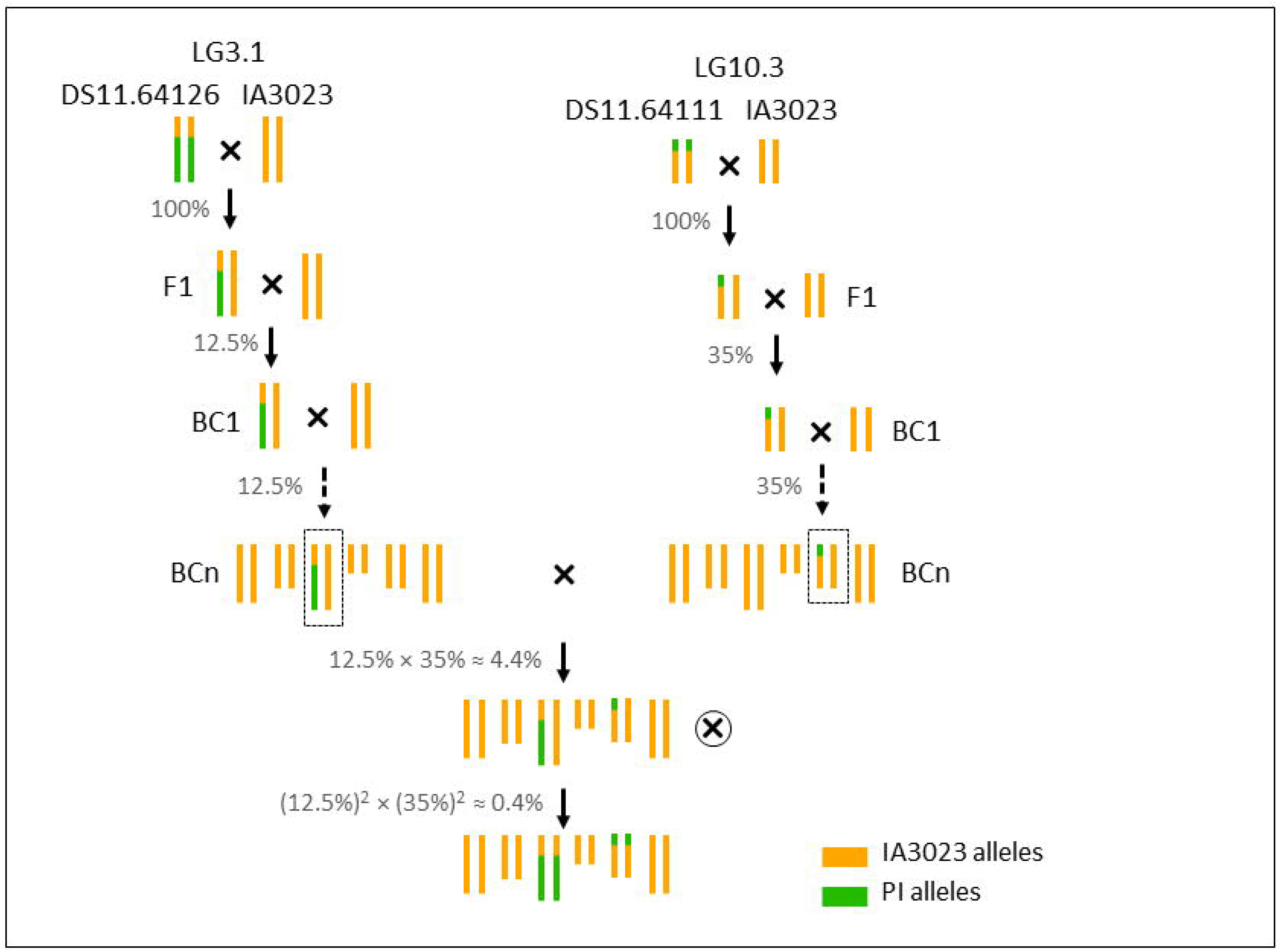
Breeding scheme for introgressing and pyramiding LGs 3.1 and 10.3 from NAM64 inbreds into IA3023.

After each round of backcrossing, 12.5% of the offspring are expected to carry one copy of LG3.1 from DS11.64126 and 35% of the offspring are expected to carry one copy of LG10.3 from DS11.64111. When selected BCn lines from two backcrossing procedures are crossed to pyramid targeted recombinations, the expected proportion of offspring carrying one copy of LG3.1 from DS11.64126 and one copy of LG10.3 from DS11.64111 is 12.5% × 35% = 4.4%. After one round of selfing, the expected proportion of offspring with the desired chromosomes is (12.5%)^2^ × (35%)^2^ = 0.4%.

## Discussion

Our results suggested that introgressing specific chromosome segments is an effective way to introduce exotic soybean germplasm into an elite cultivar. When introgression was modeled for yield in seven PI × IA3023 populations, the predicted gains ranged from 12 to 38% over the yield of IA3023. However, these gains were accompanied by an unacceptable increase of up to 14 d in maturity. This result reflected the positive correlation of 0.40 between yield and maturity in the soybean NAM populations (Diers et al. 2018). When introgression was modeled for yield while controlling maturity in the seven PI × IA3023 populations, the predicted gains remained substantial (8 to 25% over the yield of IA3023) and were accompanied by a reduction in maturity. Correlated changes in lodging, 100-seed weight, protein content, and oil content were generally small but were in the favorable direction. These results indicated that selectively assembling chromosome segments via targeted recombination may be an effective way of overcoming unfavorable correlations among traits, particularly with exotic germplasm. The PIs in this study originated from South Korea, Russian Federation, Serbia, and China (Diers et al. 2018). Our finding that germplasm from such diverse geographic origins can be used to improve a U.S. cultivar is an encouraging result for soybean breeders.

The results provided further evidence that selection in an exotic × elite cross is an ineffective way of introgressing exotic germplasm when the exotic parents are inferior to the elite parent. This inferiority was evidenced by the yield of the seven PI parents being only 59 to 81% of the yield of IA3023. This inferiority of the PIs was also reflected by the negative mean allelic effect (−52 kg/ha) for yield across PI founders in the soybean NAM study (Diers et al. 2018). Selecting for high-yielding inbreds in the seven PI × IA3023 populations led to negative or minimal gains over IA3023, with the small, positive gain of 46 kg/ha in NAM64 being accompanied by a 2 d increase in maturity (Table 2). These results were consistent with previous studies that showed the difficulty of yield improvement in exotic × elite crosses in soybean (Ininda et al. 1996, Vello et al. 1984) and maize (Hallauer and Sears 1972, Albrecht and Dudley 1987, Holland et al. 1996).

Association mapping in the soybean NAM populations had previously indicated a lack of major QTL for yield (Diers et al. 2018), and this result limited the opportunity for yield improvement through the introgression of major-effect QTL alleles from the PIs. Among a total of 23 significant yield QTLs detected in the soybean NAM study, positive yield alleles, all with small effects (≤20 kg/ha), were present in one or more PIs for only four out of the 23 QTLs (Diers et al. 2018). These few QTL with only small effects lead to small predicted gains. Consider the two positive yield alleles in PI561370 (parent of NAM50), the first on chromosome 3 with a yield effect of 11 kg/ha and the second on chromosome 11 with an effect of 12 kg/ha. If these two QTL are introgressed into IA3023, the yield gain is predicted to be only 23 kg/ha. This predicted gain is far smaller than the predicted gain of 754 kg/ha for yield with controlled maturity in NAM50 (Table 2). Our results therefore supported the effectiveness of targeted introgression of exotic germplasm even in the absence of major QTL alleles. This result is presumably due to many small-effect QTL alleles on the PI chromosome segments.

On the other hand, the large predicted gains in Table 2 assume introgression of chromosome segments on each of about 30 LGs. Introgression of up to 30 chromosome segments by backcrossing will be virtually impossible. The results showed, however, that introgressing ≤4 LGs would be sufficient in six out of the seven PI × IA3023 populations. In particular, introgressing ≤4 LGs led to a predicted yield gain (170 kg/ha) that was equal to the predicted gain in NAM03, which involved an elite × elite cross. While this gain of 170 kg/ha is much smaller than that for targeted recombination on all LGs, the goal in introgressing exotic germplasm is not necessarily to obtain the highest possible performance, but instead to widen the genetic base without decreasing the performance for key traits (Carter et al. 2004). Our results indicated that introgression of targeted recombinations on only a few LGs may achieve this goal.

We studied the usefulness of targeted recombination for exotic germplasm via a two-step approach: we first assessed the predicted gains from having 1-2 targeted recombinations per LG, then identified recombinant inbreds that carried such recombinations within each PI × IA3023 population. A recombinant inbred that had the targeted recombinations on a given LG was absent in 86% of the cases. Furthermore, some recombinant inbreds had more than two recombinations on a particular LG, and this higher number of recombinations led to a higher performance compared with the 1-2 recombinations that were modeled. These results suggested that in practice, our first step of modeling 1-2 recombinations could be omitted. Instead, the best donor recombinant inbreds can be identified for each LG, the amount of gain attributable to each LG from a donor can be predicted, the LGs can be prioritized, and a backcrossing approach for a few (e.g., ≤4) prioritized LGs with specific donors can be designed.

The backcrossing process should involve marker-assisted foreground selection for the LGs being introgressed, and marker-assisted background selection to enhance the recovery of the recurrent parent genome. This strategy relies on having an LG passed intact from a donor recombinant inbred to an offspring during backcrossing. Our analysis of LG3.1 and LG10.3 in NAM64 indicated that the probability of inheriting the desired homologue intact is reasonably high (12.5 to 35%), although this probability is expected to vary according to the size of the LG.

In NAM64, backcrossing and pyramiding LG3.1 and LG10.3 from their respective donor recombinant inbreds to IA3023 would take 3 + *n* seasons, where *n* is the number of backcross generations needed to recover the genome of IA3023 (Fig. 2). Backcrossing in soybean typically requires 3-4 generations (Kim et al. 2008, Landau-Ellis and Pantalone 2009, Kim et al. 2012). Because the donor parent already has (on average) half of the alleles in the recurrent parent, we speculate that 2-3 generations of backcrossing would be sufficient. Altogether, it would take 5-6 generations to pyramid two LGs into IA3023. Targeted introgression can be shortened through winter nurseries (Pham, Shannon and Bilyeu 2012) or CO_2_ supplemented growth chambers that enable up to five generations of soybean each year (Nagatoshi and Fujita 2018). We plan to conduct empirical experiments to assess the gains from introgressing ≤4 LGs from one of the PIs into IA3023.

## Acknowledgements

We thank Dr. Brian Diers and the soybean NAM community for conducting the NAM experiment and making the data publicly available. We thank Dr. Aaron Lorenz at University of Minnesota for generally sharing his knowledge in soybean breeding.

