## Supplementary file for "Predicted genetic gains from introgressing chromosome segments from exotic germplasm into an elite soybean cultivar"

**S1. Derivation of the probability of observing parental genotype on a linkage group during the F_5_ recombinant inbred lines (RILs)**

**Assumption**: for a targeted linkage group, the frequency of having no recombination during meiosis is *q* and the frequency of having at least one recombination during meiosis is *p* = 1 – *q*.

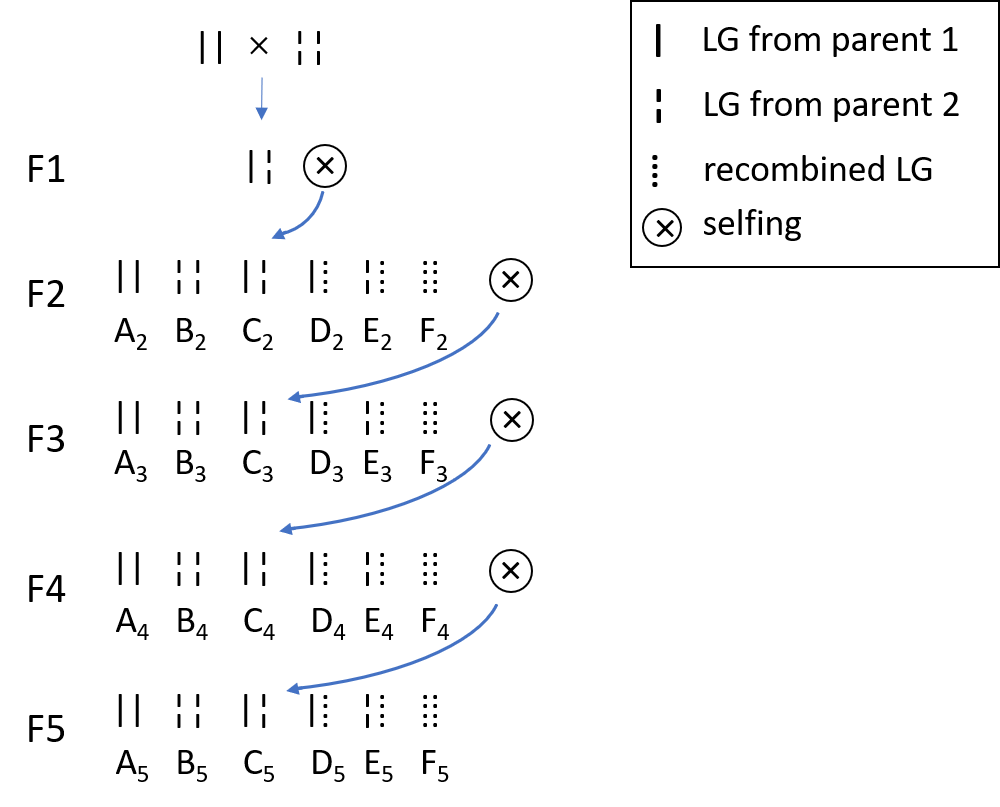

**Figure 1. breeding scheme for obtaining F_5_ RILs.** A_n_, B_n_, C_n_, D_n_, E_n_, F_n_ are the probabilities of having genotype ||, ¦¦, |¦**,** |⁞**,** ¦⁞, and ⁞⁞ on the targeted linkage group in Fn, respectively.

**Genotype distribution in F2:**

|  | A_2_ | B_2_ | C_2_ | D_2_ | E_2_ | F_2_ |
| --- | --- | --- | --- | --- | --- | --- |
| Frequency | ${\frac{1}{4}q}^{2}$ | ${\frac{1}{4}q}^{2}$ | ${\frac{1}{2}q}^{2}$ | $pq$ | $pq$ | $p^{2}$ |

**Genotype distribution in F3**

|  | A_3_ | B_3_ | C_3_ | D_3_ | E_3_ | F_3_ | Frequency in F2 |
| --- | --- | --- | --- | --- | --- | --- | --- |
| \|\| x \|\| | 1 | 0 | 0 | 0 | 0 | 0 | ${\frac{1}{4}q}^{2}$ |
| ¦¦ x ¦¦ | 0 | 1 | 0 | 0 | 0 | 0 | ${\frac{1}{4}q}^{2}$ |
| \|¦ x \|¦ | ${\frac{1}{4}q}^{2}$ | ${\frac{1}{4}q}^{2}$ | ${\frac{1}{2}q}^{2}$ | $pq$ | $pq$ | $p^{2}$ | ${\frac{1}{2}q}^{2}$ |
| \|⁞ x \|⁞ | ${\frac{1}{4}q}^{2}$ | 0 | 0 | ${\frac{1}{2}q}^{2}+pq$ | 0 | ${\frac{1}{4}q}^{2}+pq+p^{2}$ | $pq$ |
| ¦⁞ x ¦⁞ | 0 | ${\frac{1}{4}q}^{2}$ | 0 | 0 | ${\frac{1}{2}q}^{2}+pq$ | ${\frac{1}{4}q}^{2}+pq+p^{2}$ | $pq$ |
| ⁞⁞ x ⁞⁞ | 0 | 0 | 0 | 0 | 0 | 1 | $p^{2}$ |

$$A_{3}={\left( 1 \right)A}_{2}+\left( \frac{1}{4}q^{2} \right)C_{2}+\left( \frac{1}{4}q^{2} \right)D_{2}$$

$$=\frac{1}{4}q^{2}+ \left( \frac{1}{4}q^{2} \right)\left( \frac{1}{2}q^{2} \right)+\left( \frac{1}{4}q^{2} \right)pq$$

$$=\frac{q^{2}}{8}(q^{2}+2pq+2)$$

$$B_{3}=\left( 1 \right)B_{2}+\left( \frac{1}{4}q^{2} \right)C_{2}+(\frac{1}{4}q^{2})E_{2}$$

$$=\frac{1}{4}q^{2}+ \left( \frac{1}{4}q^{2} \right)(\frac{1}{2}q^{2})+\left( \frac{1}{4}q^{2} \right)pq$$

$$= \frac{q^{2}}{8}(q^{2}+2pq+2)$$

$$C_{3}=\left( \frac{1}{2}q^{2} \right)C_{2}$$

$$= \frac{1}{4}q^{4}$$

$$D_{3}=\left( pq \right)C_{2}+\left( {\frac{1}{2}q}^{2}+pq \right)D_{2}$$

$$=pq\left( \frac{1}{2}q^{2} \right)+\left( {\frac{1}{2}q}^{2}+pq \right)pq$$

$$= pq^{2}$$

$$E_{3}=\left( pq \right)C_{2}+\left( {\frac{1}{2}q}^{2}+pq \right)E_{2}$$

$$=pq\left( \frac{1}{2}q^{2} \right)+\left( {\frac{1}{2}q}^{2}+pq \right)pq$$

$$= pq^{2}$$

$$F_{3}=\left( p^{2} \right)C_{2}+\left( {\frac{1}{4}q}^{2}+pq+p^{2} \right)\left( D_{2}+E_{2} \right)+\left( 1 \right)F_{2}$$

$$=p^{2}\left( \frac{1}{2}q^{2} \right)+\left( {\frac{1}{4}q}^{2}+pq+p^{2} \right)2pq+ p^{2}$$

$$=pq\left( 2p^{2}+\frac{5}{2}pq+\frac{1}{2}q^{2} \right)+p^{2}$$

**Generalized equations from F_n_ to F_n+1_ (n >= 2)**

$$A_{n+1}={\left( 1 \right)A}_{n}+\left( \frac{1}{4}q^{2} \right)C_{n}+\left( \frac{1}{4}q^{2} \right)D_{n}$$

$$B_{n+1}=\left( 1 \right)B_{n}+\left( \frac{1}{4}q^{2} \right)C_{n}+(\frac{1}{4}q^{2})E_{n}$$

$$C_{n+1}=\left( \frac{1}{2}q^{2} \right)C_{n}$$

$$D_{n+1}=\left( pq \right)C_{n}+\left( {\frac{1}{2}q}^{2}+pq \right)D_{n}$$

$$E_{n}=\left( pq \right)C_{n}+\left( {\frac{1}{2}q}^{2}+pq \right)E_{n}$$

$$F_{n}=\left( p^{2} \right)C_{n}+\left( {\frac{1}{4}q}^{2}+pq+p^{2} \right)\left( D_{n}+E_{n} \right)+\left( 1 \right)F_{n}$$
